# Oddball evoked deviant responses reflect complex context dependent expectations in mouse V1

**DOI:** 10.1101/2024.09.30.615789

**Authors:** Scott G. Knudstrup, Catalina Martinez Reyes, Cambria M. Jensen, Rachel W. Schecter, Mac Kenzie Frank, Jeffrey P. Gavornik

## Abstract

Evoked responses in the mouse primary visual cortex can be modulated by the temporal context in which visual inputs are presented. Oddball stimuli embedded in a sequence of regularly repeated visual elements have been shown to drive relatively large deviant responses, a finding that is generally consistent with the theory that cortical circuits implement a form of predictive coding. These results can be confounded by short-term adaptation effects, however, that make interpretation difficult. Here we use various forms of the oddball paradigm to disentangle temporal and ordinal components of the deviant response, showing that it is a complex phenomenon affected by temporal structure, ordinal expectation, and event frequency. Specifically, we use visually evoked potentials to show that deviant responses occur over a large range of time, lead to long-term plasticity in some cases, cannot be explained by a simple adaptation model, scale with predictability, and are modulated by violations of both first and second-order sequential expectations.

**Significance Statement:** Visual experience and temporal context can modulate evoked responses in mouse V1. There remains disagreement about whether this reflects predictive coding in visual circuits and if visual mismatched negativity, which has important cross-over implications for human clinical work, constitutes evidence supporting this theory or simple neural adaptation. This work strongly supports the former interpretation by demonstrating complex experience-dependent deviant responses that cannot be easily explained by a simple adaptation model. We use statistically rigorous analysis of the local field potential to show that oddball evoked deviance signals reflect relative timing, event frequency, 1^st^ and 2^nd^ order sequence expectations and scale as a function of event probability.

## Introduction

Unexpected “oddball” elements inserted in otherwise predictable sequences can drive large transient responses in sensory cortex that may represent evidence of predictive coding, a model of brain function suggesting that neural responses are inversely proportional to their predictability (Rao and Ballard, 1999; Friston, 2005; Keller and Mrsic-Flogel, 2018). While the most well-known and studied form of this phenomenon is Mismatched Negativity (MMN) in the auditory system (Nelken and Ulanovsky, 2007; Wacongne et al., 2012; Fitzgerald and Todd, 2020), a similar phenomenon also exists in the visual system (Pazo-Alvarez et al., 2003; Stefanics et al., 2014, 2015). Mice have emerged as an important model in which to study the mechanistic basis of this signal (Kremláček et al., 2016; Ross and Hamm, 2020), which is abnormal in schizophrenia, autisms spectrum disorder, and other conditions (Shelley et al., 1991; Neuhoff et al., 2012; Hirt et al., 2019; Lassen et al., 2022; Molnár et al., 2024) but difficult to study in humans for obvious reasons. Several recent works in mouse have used genetically targeted fluorescent indicators and a 2-photon (2P) microscope to map the circuits involved with oddball-evoked responses in the primary visual cortex (V1)(Hamm and Yuste, 2016; Hamm et al., 2021; Homann et al., 2022; Gillon et al., 2024) and frontal areas (Van Derveer et al., 2023) using variations of a visual oddball paradigm. One work in particular (Homann et al., 2022) expanded the scope of this phenomenon by using randomly positioned Gabor patches over a gray background to create multi-element sequence structures and showed that novel oddball stimuli drive excess calcium signal in a large population of L2/3 neurons. Substituting a familiar but out-of-order element did not produce an obvious oddball response, leading the authors to propose that neuronal-adaptation is the primary mechanism driving elevated responses to unexpected, novel images.

This adaptation-based model is somewhat at odds with our previous work showing that ordinal sequence violations (created by re-ordering elements of a known sequence rather than substituting a novel element) and temporal violation (created by showing an expected element at an unexpected time) can drive significant differences in V1 activity after exposing mice to a single training sequence for several days. These effects are visible in visually evoked potentials (VEPs) calculated from local field potential (LFP) recordings, in spiking neuron activity recorded with extracellular electrodes, and in 2-photon Ca^2+^ imaging (Gavornik and Bear, 2014; Sidorov et al., 2020; Finnie et al., 2021; Price et al., 2023; Knudstrup et al., 2024a) and have lead us to conclude that experience-dependent plasticity allows V1 to learn complex predictive spatiotemporal codes (Price and Gavornik, 2022).

Both single-day oddball and multiday sequence learning experiments reveal the capacity of V1 to encode associations between spatiotemporal stimuli and differentially signal expectation violations. The visually evoked activity during sequence learning training that drives long-term plasticity across days is presumptively similar to evoked activity during oddball experiments. It is important to clarify whether oddball evoked deviant responses within a single session occur following complex spatiotemporal expectation violations seen after multi-day induction, especially since previously published experiments differ in time scale (single-day or multi-day), stimulus design (visual stimulus complexity), violation type (familiarity, ordinal, or temporal), and recording methodology and location (two-photon in L2/3 vs electrophysiology in L4). In this study we conducted a set of single-session oddball experiments to characterize deviant responses driven by different stimuli and violation types. We then determined the range of stimulus durations over which oddball responses occur and showed that temporal structure is important, deviant responses do not occur based on temporal expectation violations. Finally, we showed that deviant responses scale as a function of oddball event frequency rather than the inter-oddball interval, that deviant responses are generally larger when evoked by less-predictable oddball events, and that they can occur following higher-order expectation violations.

## Methods and Materials

*Animals.* Male and Female C57BL/6 (Charles River Laboratories), non-carrier C57BL/6J-Tg(Thy1-GCaMP6f)GP5.17Dkim/J (The Jackson Laboratory, 025393), non-carrier B6.SJL-Slc6a3tm1.1(cre)Bkmn/J (The Jackson Laboratory, 006660), and non-carrier B6.Cg-Tg(Slc32a1-COP4*H134R/EYFP)8Gfng/J (The Jackson Laboratory, 014548) mice were used. Mice were group housed with littermates (max. five mice per cage) on a 12-h light/dark cycle and provided food and water ad libitum. All experiments were performed during the light-cycle. All procedures were approved by the Institutional Animal Care and Use Committee of Boston University.

### Electrode implantation

Mice were anesthetized with an intraperitoneal injection of 50 mg/kg ketamine and 10 mg/kg xylazine. To facilitate head restraint, a steel headpost was affixed to the skull anterior to bregma using cyanoacrylate glue. Small (<0.5 mm) burr holes were drilled over primary visual cortex (3 mm lateral and 0.4 mm anterior to lambda) and tungsten microelectrodes (FHC) were placed 450 μm below the cortical surface. Proper placement was confirmed based on visual responsiveness and VEP waveform morphology or via postmortem histology. Reference electrodes (silver wire, A-M systems) were placed below dura over left parietal cortex. All electrodes were rigidly secured to the skull using cyanoacrylate glue. Dental cement was used to enclose exposed skull and electrodes in a protective head cap. Buprenex (0.1 mg/kg) was injected subcutaneously for postoperative pain amelioration. Surgery was performed between postnatal day 45-90. Mice were monitored for signs of infection and allowed at least 24 h of recovery before habituation to the recording and restraint apparatus and were excluded from experiments only in the event of unsuccessful electrode implantation (lack of visual response, incorrect depth, poor grounding, etc.). To maximize yield, electrodes were implanted bilaterally with the right electrode intended as a spare in case the left electrode failed. In most cases the left hemisphere electrode produced clear VEPs and was used for data recording and analysis.

### Data collection

All data was amplified and digitized using the digital Recorder-64 system (Plexon). Local field potentials (LFPs) were recorded with 1-kHz sampling and a 200-Hz low-pass filter. Data was extracted from the binary storage files and analysed using custom software written in C++ and MATLAB, all of which is available for download at https://gavorniklab.bu.edu/supplemental-materials. Reported n values reflect the number of mice included in each study. In some cases individual mice were used in multiple experiments with different sequences (e.g. sequences composed of different visual elements) presented in different recording sessions.

### Visual stimulation

Visual stimuli were generated and displayed using MATLAB with the PsychToolbox extension (Brainard, 1997), with custom software (https://github.com/jeffgavornik/VEPStimulusSuite) used to control timing and hardware signals. Stimuli were displayed on a 22-inch LED monitor (1920 x 1080 pixels, refresh rate 60 Hz) positioned 25 cm directly in front of the mouse. Oriented sinusoidal gratings (OSG, 0.5 cycles per degree) were separated by a minimum of 30 degrees and shown at full contrast. Multi Random Gabor (MRG) stimuli were composed of superpositions of Gabor patches. For each image, 100 Gabor functions were chosen with random location, orientation, and phase, and either ON- or OFF-polarity. The spatial extent for each Gabor function was randomly selected to be between 10 and 20 degrees, matching the receptive field sizes of V1 in mice. Gabor patches were linearly superimposed with saturation at 100% contrast. All stimuli spanned the full range of monitor display values between black and white, with gamma correction to ensure a linear gradient and constant total luminance.

### Statistics

Most statistics were conducted using the nonparametric bootstrap for multilevel data (Saravanan et al., 2020). For each iteration of the bootstrap, a random subset of mice was selected (with replacement). Event-locked LFPs for all standard trials and all selected mice were combined into a single matrix (trials x time), and the same was done for deviant trials. For each of these two matrices, a random selection of rows/trials was drawn (with replacement) equal to the number of rows/trials in the matrix. These selections were then averaged over row/trial, yielding a single mean standard trace and a mean deviant trace. This was repeated 1000 times to generate a distribution of standard and deviant mean traces. 95% confidence intervals were generated by finding the 2.5^th^ and 97.5^th^ percentile of the bootstrapped traces for each time point individually. Plots of bootstrapped standard and deviant traces display the median bootstrap estimate as solid lines with the with 95% percentile confidence intervals in shaded regions. Difference distributions were computed by taking the element-wise difference between the bootstrapped mean traces (standard – deviant). Plots of bootstrapped difference traces show the inner 95% and distribution median in shaded and solid lines, respectively. Time points that do not contain zero within the 95% confidence interval indicate where standard and deviant distributions are significantly different, and we indicate these time points with cyan lines along the x-axis. When multiple comparisons were made within a single experiment, statistics were computed after scaling confidence intervals using a conservative Bonferroni correction.

The oddball score (OBS) is defined as the time averaged absolute difference between standard and deviant traces over a given window with units of µV/ms. With a sampling rate of 1000 Hz, the oddball score is computed as 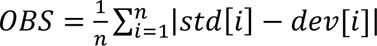, where *std* and *dev* are traces consisting of n samples each. Oddball score distributions were generated by computing the OBS bootstrapped standard and deviant trace estimates using the nonparametric hierarchical bootstrap procedure described above. We also fit a linear model based on OBS scores computed directly (i.e., non-bootstrapped) by first calculating the mean standard and deviant traces for a given mouse and using those to compute the OBS. In this case, each mouse generates a single OBS value. Model fitting on oddball scores was performed with the Python library statsmodels.

## Results

### Visual oddballs drive significant deviant responses with different stimulus types

We first investigated whether sequence-evoked VEPs recorded in layer four (L4) of binocular V1 exhibit deviant responses, and the dependence on stimulus type, by exposing head-fixed mice (Fig. 1A) to repeated presentations of a four-image standard sequence (Std, ABCD, Fig. 1B). Every tenth presentation, a deviant sequence was created by replacing element B with the novel oddball element E (Dev, AECD). All mice (n=14) were tested with sequences consisting of oriented sinusoidal gratings (OSG, as used in previous multi-day sequence learning experiments) and randomly placed Gabor patches (RPG, similar to Homann et al. 2022). In both cases, sequence elements persisted for 200 ms, and the animals passively viewed a total of 500 sequences (450 Std, 50 Dev) in a single recording session. LFPs were measured during visual stimulation, and we used a nonparametric hierarchical bootstrap approach (Saravanan et al., 2020) to estimate VEPs and perform statistical comparisons (Fig. 1C). Briefly, bootstrap estimates of Std and Dev VEPs (N_BootStrap_ = 1000) were calculated as the population mean after using random resampling with replacement to select both animals and trials. Plotted VEPs (Fig. 1D) represent the median timeseries from the resulting distribution with shaded regions showing 95% confidence intervals unless otherwise noted.

**Fig. 1.**
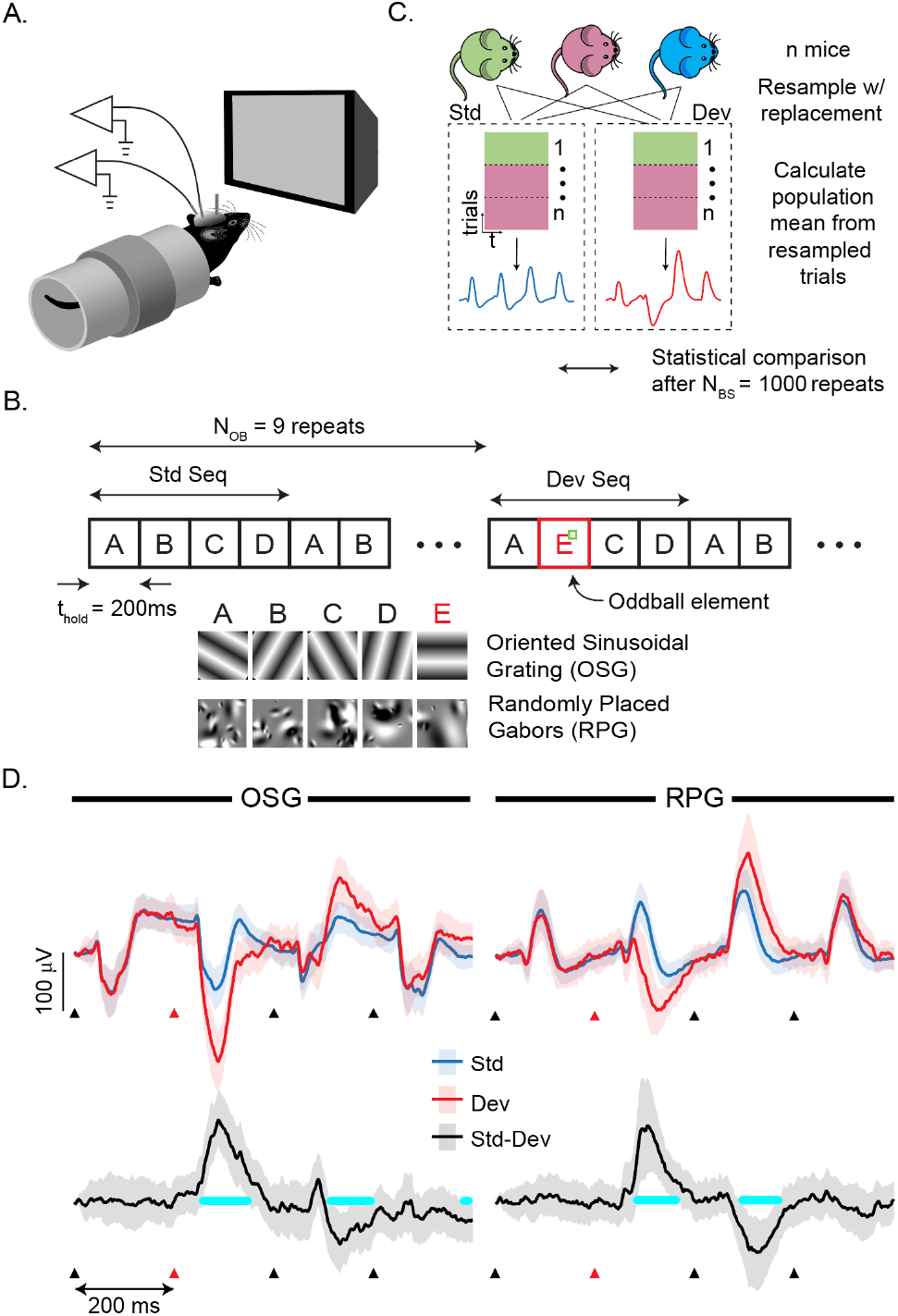
Visual oddballs produce deviant LFP responses. **(A)** Mice with LFP recording electrodes chronically implanted in binocular V1 L4 were head-fixed in front of a computer monitor. **(B)** Visual stimulation consisted of repeated presentations of a standard sequence, ABCD, with an oddball element E replacing B every 10^th^ presentation. Each element was held on the screen for 200 ms and sequences were presented 500 times (450 Std and 50 Dev sequences per session). Individual elements consisted of either an oriented sinusoidal grating (OSG) or randomly placed Gabor patches (RPG). (**C)** A hierarchical bootstrap approach was used for nonparametric statistical comparisons. Briefly, we made 1000 estimates of Std and Dev responses by calculating the mean VEP after resampling at the mouse and trial levels and used non-parametric statistics to compare the resulting distributions (one bootstrap exemplar diagramed here, see methods). **(D)** Both the OSG and RPG sequences drove stereotypical VEPs in response to the Std sequence (blue). In both cases, the Dev response (red) differed markedly when the oddball element E was shown. In this and subsequent plots, solid lines represent the median VEP calculated from bootstrapped means, shaded regions show 95% confidence intervals unless otherwise indicated, and triangles mark sequence element onset times (oddball locations in red). Std and Dev responses are significantly different when the confidence intervals around the difference between the two (Std-Dev, black trace bottom) does not include zero. Significantly different time points are marked by cyan. For both stimuli, the Dev response is significantly different from Std for a period of time starting immediately after the oddball presentation and extending into the next sequence element C. Note that element C occurs in exactly the same location within the Std and Dev sequences.

Both OSG and RPG stimuli drove reliable VEPs, though with different characteristic waveform shapes (Fig. 1D). In both cases, inclusion of a novel element drove a Dev response that was significantly different from the Std response at the 95% confidence level (equivalent to a type-1error rate of α=0.05, identified by periods of time when the 95% confidence intervals of the Std-Dev distribution do not include zero) (Saravanan et al., 2020; Hayden et al., 2021, 2023). Despite the differences in VEP shapes evoked by the two stimuli classes, the delta between Std and Dev looks very similar. In both cases, the period of significant difference appeared rapidly and persisted well into the subsequent element C response despite the fact that element C occurred at the same ordinal and temporal location within the sequence as in the Std case. On average, the significant Dev response emerged slightly faster after element E for OSG relative to RPG, 60 ms vs 82 ms, but did last quite as long, 367 ms vs 391 ms.

### Activity dependent plasticity is influenced by the temporal structure of visual stimuli

Our previous work focused on characterizing how spatiotemporal predictions emerge in V1 circuits following multiple days of sequence exposure. While superficially similar, the sequence structure used in these studies varied from most oddball paradigms in that individual sequence presentations were separated by a relatively long period of uniform gray screen (in most of our studies the intersequence period was 1.5 secs, approximately twice as long as the sequence itself). While the plasticity resulting from our multi-day paradigm is readily visible in the LFP (Gavornik and Bear, 2014; Sidorov et al., 2020; Finnie et al., 2021; Sarkar et al., 2024) and at single cell levels (Price et al., 2023; Knudstrup et al., 2024a), evidence for long-term plasticity resulting from a continuous oddball paradigm is less clear. To test whether inclusion of an interstimulus gray screen affects plasticity levels, we exposed groups of mice to fixed sequences (ABCD) presented continuously without gray screen 200 times per day for 5 days. Comparing VEPs produced by this sequence on days 1 and 5 (Fig. 2, left) demonstrates that while there is a clear change in the negative-going responses to each sequence elements, the magnitude of this change is small (on order of 10s of μVs) compared to that reported in previous works (order of 100s of μVs). Interestingly, the downward going response after element onset corresponding to the arrival of retinofugal inputs at V1 L4 is not significantly different between exposure days 1 and 5, with a significant difference emerging only later during the period of sustained input suggesting minimal plasticity of thalamocortical synapses in V1.

**Fig. 2.**
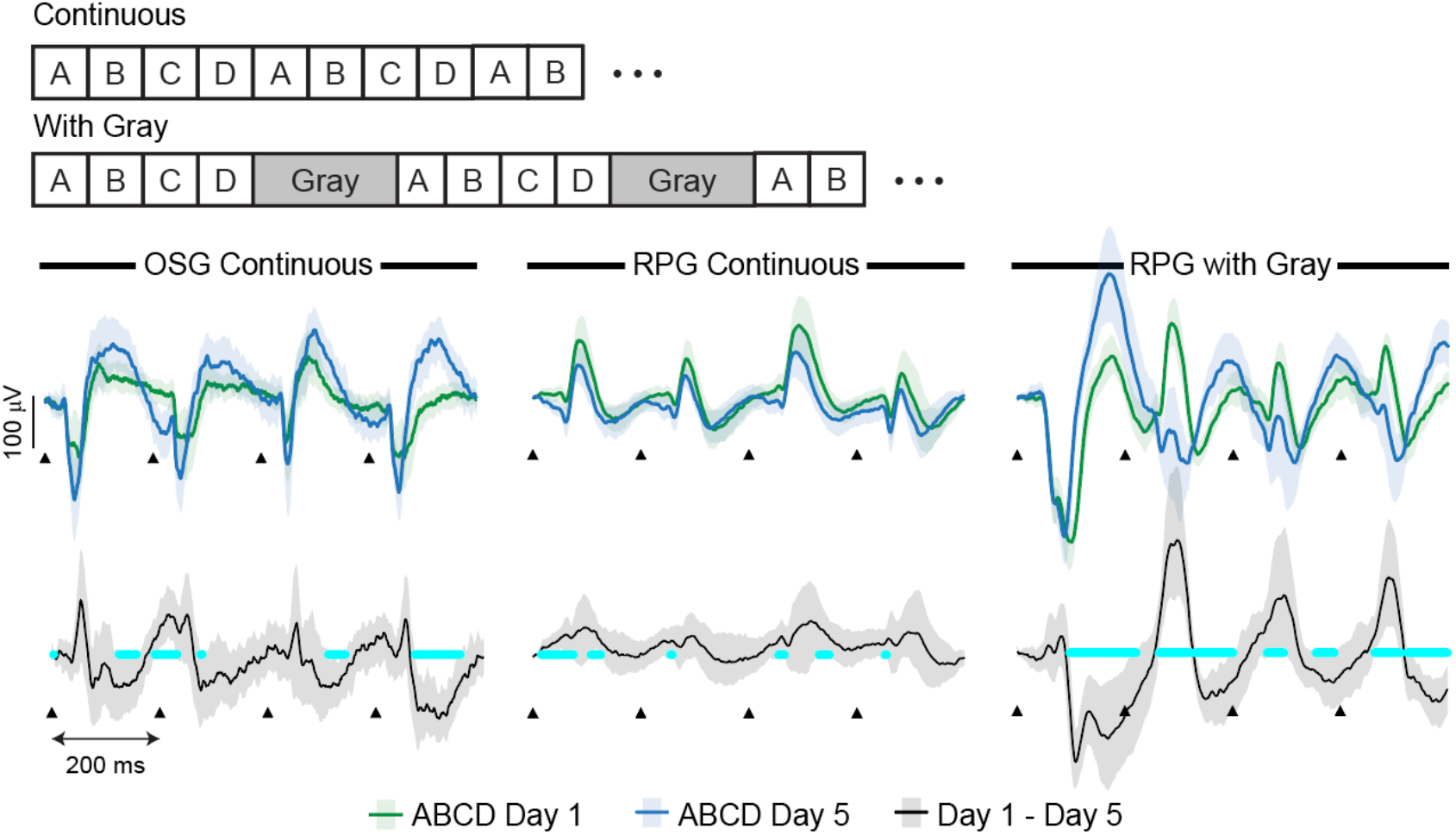
Continuous RPG sequences drive minimal plasticity. Mice viewed a fixed sequence, ABCD, 200 times per day for five days. Sequences were composed of oriented sinusoidal gratings (OSG, n=8), randomly placed Gaussians (RPG, n=5), or RPG with 1.5 sec of gray separating each sequence presentation (n=6). The top row shows averaged evoked potentials recorded in response to the sequence on Days 1 (green) and 5 (blue) with 99% confidence intervals and the bottom shows the difference between days. While there was a significant difference between Day 1 and 5 responses in all groups (cyan bars indicate periods of significant difference), RPG resulted in minimal changes to the size and shape of evoked responses. Dramatically more plasticity occurred when RPG sequence presentations were separated with a gray screen. OSG-evoked plasticity is also smaller than in previously published experiments (e.g. Gavornik and Bear 2014) where sequences were also separated with a gray screen.

We next tested whether continuous RPG stimuli also produce plasticity effecting evoked potential dynamics. Surprisingly, there was comparatively little change in VEP waveforms (Fig. 2, center) compared to that seen with the OSG stimulus and the magnitude of change increased considerably when the same RPG sequence was shown with gray (Fig. 2, right). Since RPG images drive large easily detectable deviant responses, result in relatively minimal long-term plasticity, and allow a space of unique images that is much greater than oriented gratings (practically restricted to 6 unique orientations between 0 and 180°), we chose to use RPG sequences for the remainder of our studies on the oddball phenomenon.

### Oddball responses are mediated by, but do not predict, temporal structure

Our next aim was to determine the temporal range over which oddball responses can be elicited by familiarity violations. Toward this end, and to expand upon previous works that used stimulus hold times in the 150-300 ms range, we repeated the familiarity violation protocol from the previous experiment (i.e., ABCD x 9 followed by AECD) under different element timing conditions. In separate sessions, mice were shown visual sequences with element times of 75 ms (n=15), 500 ms (n=15), 1000 ms (n=19), and 2000 ms (n=12). We again used the nonparametric hierarchical bootstrap to generate distributions of mean VEPs for each group (ABCD and AECD) for each timing condition.

We find that the responses to novel oddball elements were significantly different from those elicited by the standard element in all cases (Fig. 3A-C) except for 2000 ms element timing condition (Fig. 3D). As before, the significant deviation emerged rapidly after oddball element onset. The overall magnitude of the deviant response decreases with element time (compare the black Std-Dev trace in panels A and B, which are plotted on the same vertical scale) and is characterized by a deviant response to the oddball which is relatively negative compared with the Std response. For very fast times (75 ms) the period of significant difference spans adjacent sequence elements. For longer times (200 ms in Fig 1., 500 ms in Fig. 3), the element following the oddball still generates a significantly different response than in the standard condition though the period of significant difference is no longer contiguous with the initial deviant response. Note that the repeat time between expected elements (calculated as the duration between element onsets) for these cases is 0.3, 2, 4, and 8 seconds for t_hold_ = 0.075, 0.5, 1.0 and 2.0 seconds, respectively. The approximate inter-oddball intervals for the respective hold times are 3.5, 20, 40, and 80 seconds.

**Fig. 3.**
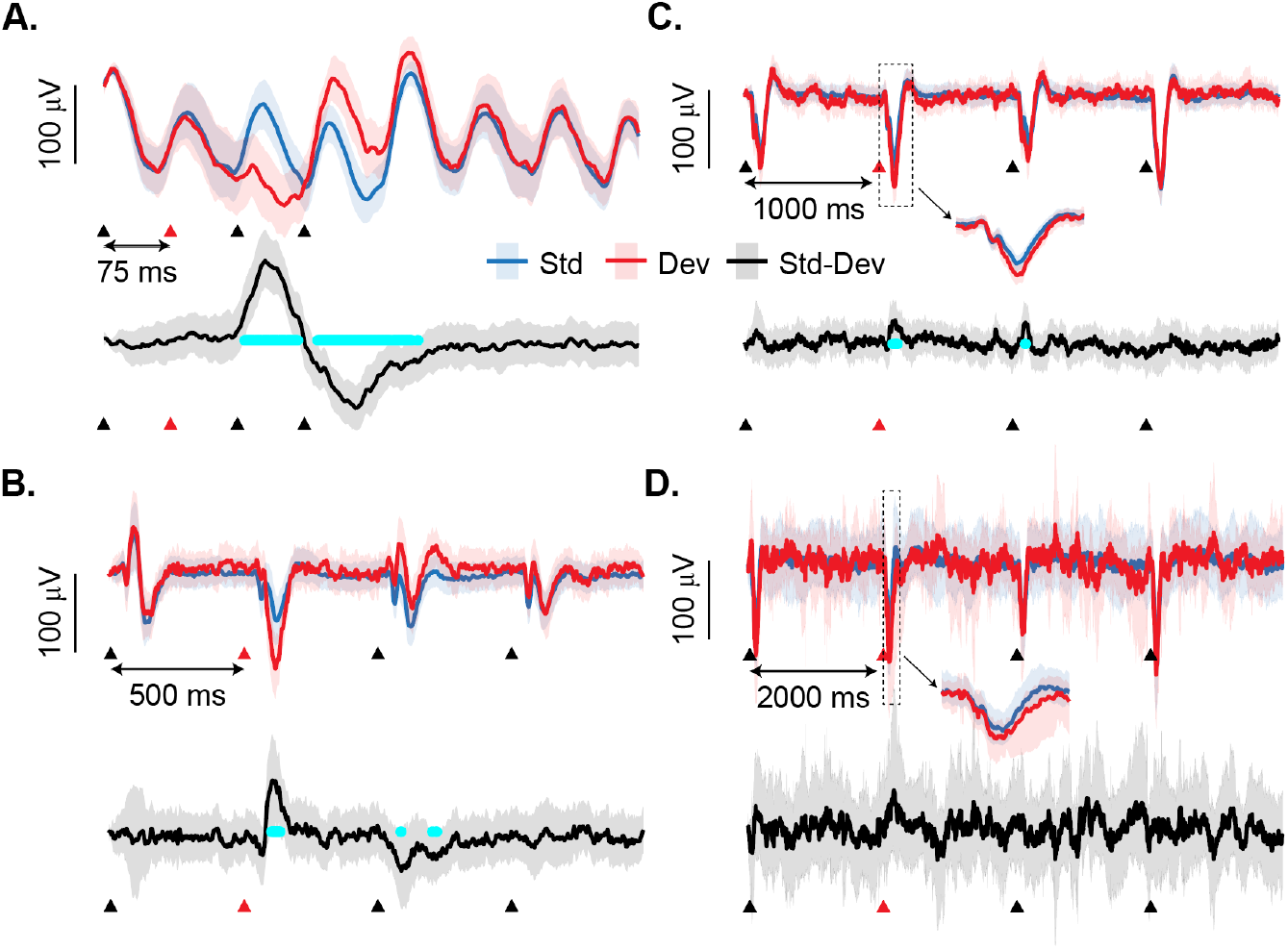
V1 signals familiarity violations over a wide range of stimulus times. Mice viewed a Std RPG sequence 9 times for every Dev sequence. In separate recording sessions, mice were shown different sequences with element durations (thold) of 75, 500, 1000, and 2000 ms (n=12 for thold=2000, n=15 for others). (A-C) Statistically significant deviant responses extending across multiple elements were seen for 75, 500, and 1000 ms though the magnitude of this effect decreased with element duration. (D) The deviant response at 2000 ms looked similar to that seen at 1000 ms (insets show 250 ms of activity evoked by the second element), but this was not statistically significant. The latency until significance also increased with thold, with values of approximately 80, 107, and 108 ms for 75, 500, and 1000 ms hold times. The time between oddball presentations was 3, 20, 40 and 80 s for thold = 75, 500, 1000, and 2000 ms.

Having established that the temporal structure of a sequence modulates multi-day plasticity and the deviant response to oddball stimuli within a single recording session, we next tested whether deviant responses require a fixed temporal structure using an oddball paradigm where each element had a unique hold time (Fig. 4A). An oddball element still generates a significant deviance response when inserted into a sequence ABCD with hold times of 200, 300, 350 and 250 ms for elements A, B, C, and D (Fig. 4A) though the response is small compared to previous experiments with a single hold time in this same range (Fig. 1D) and does not persist into the element C response. When the experiment was repeated with the hold time for every element presentation drawn from a uniform random distribution between 150 and 300 ms, the oddball element creates a deviant response that is trending towards that seen in previous experiments (e.g. relatively negative compared to the standard response), but this is no longer significant (Fig. 4B). We conclude from these experiments that the deviant responses is sensitive to the regularity of visual stimuli and can disappear completely when sequence element timing is randomized even though the expected hold times and total sequence time, on average in this case 225 ms and 900 ms respectively, is well within the temporal window where significant deviant responses were seen previously.

**Fig. 4.**
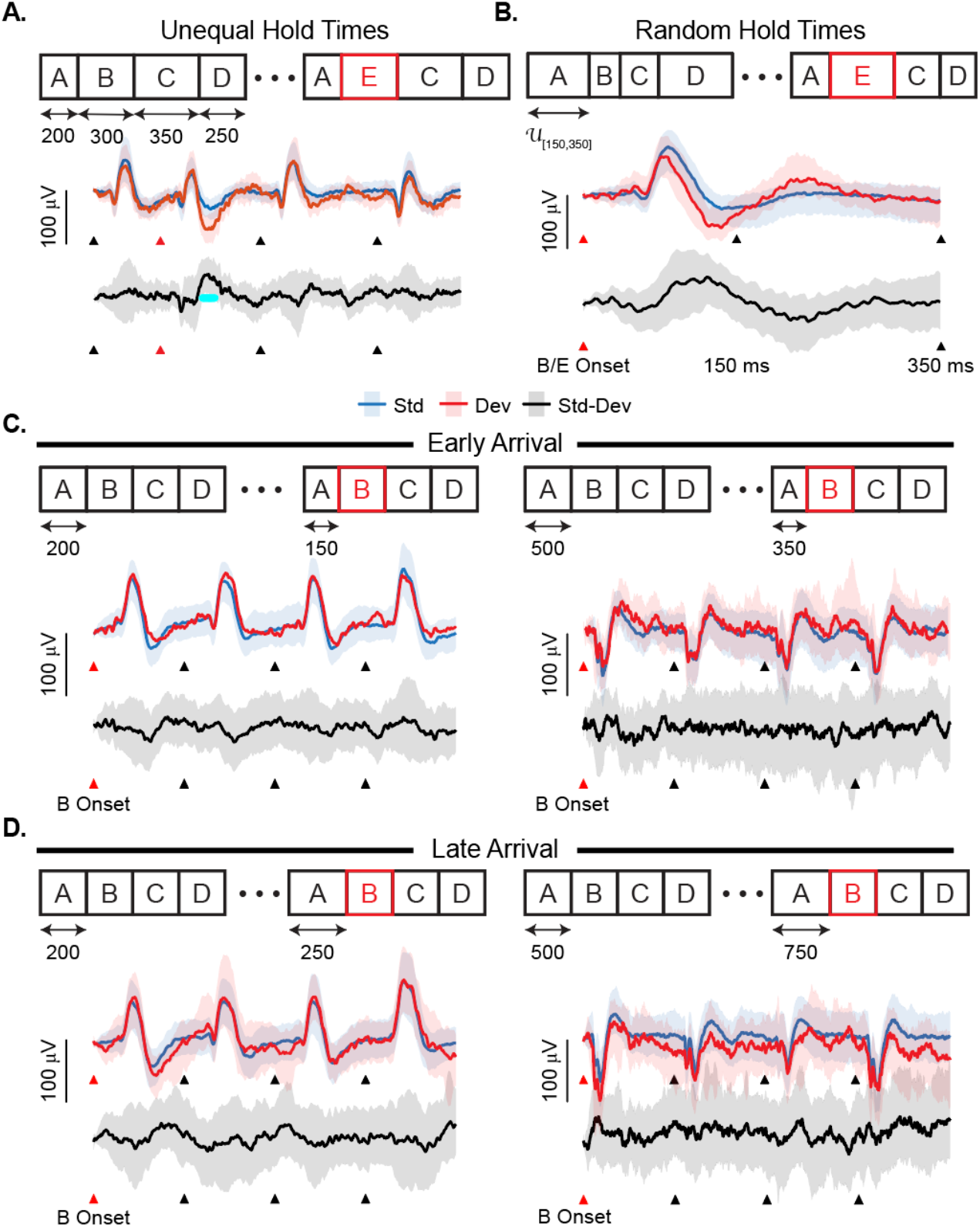
Deviant responses reflect sequence timing but do not explicitly indicate temporal expectation violations. (A) Mice were presented with an RPG sequence ABCD where each element had a different hold time (250, 300, 350, and 250 ms for A, B, C, and D). An oddball element E, which replaced B every 10^th^ presentation, drove a significantly deviant response. When compared with a fixed sequence (Fig. 1D) the deviant response was relatively late and of brief duration. (B) An oddball element did not produce a significantly different response when elements were presented with random timing (thold was selected randomly for each presentation from a uniform distribution between 150 and 350 ms). This plot shows the response aligned to the onset of elements B and E (red triangles, this alignment is required due to the varying time of preceding elements) with markers indicating the range over which subsequent elements would appear. (C-D) Creating a “temporal oddball” by presenting expected element B either early or late did not produce significant deviant responses, as shown by responses aligned to the 2^nd^ element for hold times of 200 (left, +/- 50 ms) or 500 ms (right, +/- 150 ms).

We next tested whether or not visual responses are modulated by violations of established temporal relationships by showing expected sequence elements at unexpected times. We presented mice with repeated presentations of the sequence ABCD as before, but element B was presented early (Fig. 4C) or late (Fig. 4D) by changing the hold time of element A in every 10^th^ presentation. In our first round of experiments, we used a standard hold time of 200 ms and presented B 50 ms early or late by holding element A for either 150 or 250 ms. We then repeated this experiment with a standard hold time of 500 ms, showing B 150 ms early or late by holding A for either 350 or 650 ms. In neither case did the “temporal oddball” drive a deviant response compared to the standard case. We conclude from this result that V1 responses to visual oddballs do not explicitly represent temporal expectations violations.

### Deviant responses cannot be explained by simple adaptation

Neural responses undergo adaptation following repeated presentations of the same sensory stimuli. To determine whether this effect could be responsible for mediating deviant responses to oddball stimuli, we generated 20 unique RPG elements and showed them with a 200 ms hold time to mice in a random order with a uniform probability of transitioning between elements (e.g. the probability of and A-B transition was equal to an A-C, A-D etc. transition). Self-transitions (e.g. A-A) were not allowed. Each element presentation was characterized based on how recently the same element had been seen and VEPs were estimated using our hierarchical bootstrap approach based on this offset rather than element designation. So, for example, the 2^nd^ presentation of A in A-F-A would be grouped with the second presentation of G in G-B-G and classified as a having a single element delta, or 200 ms, separating presentations. VEP estimates, shown in figure 5A with 99.5% confidence intervals to allow multiple comparisons, show that while the initial downward-going component of the VEP is the same for all offsets, the later positive-going component is largest for the 200 ms interval (red line). Pair-wise comparisons (e.g. 200ms – 400 ms) show that the difference as a function of offset is only significant for short timescales (Fig. 5B, right). To quantify the size of the effect, we calculated an “Oddball Score” (OBS) as the time-averaged absolute value of the difference between conditions being compared over the duration of the VEP (e.g. an OBS of 6 would mean that two VEPs had an average difference of 6 µV per ms over the 200 ms analysis window, see methods for details). As shown in Fig. 5B, left, the bootstrapped distribution of OBS was largest for short offset times and approximately equal for longer comparisons. Similar results with a hold-time of 400 ms do not result in any significant differences as a function of offset using the same method. These results suggest that adaptation has a significant effect on VEP shape only when the same stimulus is seen repeatedly over short time scales on the range of approximately 200-400 ms. Note that element-repeat times in previous experiments are outside of this range in all cases except the 75 ms hold time (Fig. 3A).

**Fig. 5.**
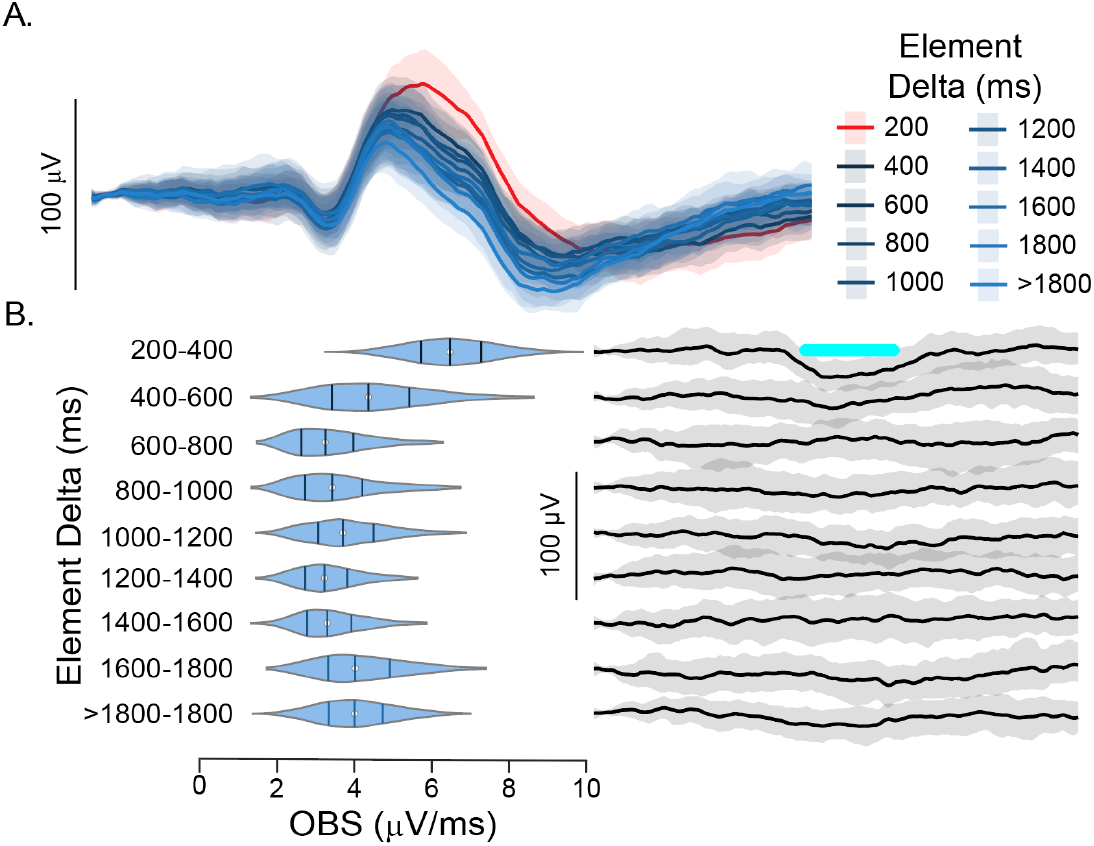
Short term adaptation significantly effects VEP magnitude over a short time. 20 RPG elements were presented 200 times each in a random order with a 200 ms hold time and an equal of probability transitioning between elements (self-transitions, e.g. A-A, were not allowed). Evoked potentials were sorted based on how recently the same element had been seen and bootstrap VEP estimates were calculated with all elements grouped together. (A) Bootstrapped VEPs as a function of Element Delta, where Element Delta represents the amount of time between the end of an element presentation and its next onset (A-B-A = 200 ms, A-B-C-A = 400 ms, etc.). Plots display median bootstrapped means (solid lines) with 99.5% confidence intervals (shaded regions, statistics calculated with Bonferroni corrected levels). The positive-going component of the response was noticeably larger (red) when repeats were separated by a single element (e.g. A-B-A) but stabilized as the Element Delta increased. (A) For each bootstrap, the oddball score (OBS) was calculated based on the difference between Element Delta population responses for each neighboring pairing (e.g., 200 ms vs 400 ms, 400 ms vs 600 ms, etc.) up to 10 element deltas. The OBS was largest when comparing the 200 and 400 ms VEPs (violin plots, left), which corresponded to a significant difference between the VEPs (delta plots, right). Violins show the estimated density of bootstrapped distributions with solid lines marking quartiles. The OBS was smaller for all other comparisons, none of which showed a significant difference between VEPs, demonstrating that adaptation has a significant effect on the VEP only when elements are repeated within a brief window.

Having identified adaptation as a clear confound for short timescales, we next set out to disentangle recency effects from oddball frequency and ordinal structure. To do so, we first constructed sequences with equal inter-deviant intervals by doubling the hold time while halving oddball frequency (e.g. 200 ms with an oddball every 10^th^ sequence presentation vs. 400 ms with an oddball every 5^th^ sequence presentation, Fig. 6A). If the deviant response created by an oddball stimulus is simply a consequence of recency or total time spent viewing each sequence element, we would expect the effect to be approximately the same for both hold times. Instead, bootstrap estimates of the OBS are larger when the oddball is presented less often but with the same amount of time for inter-deviant intervals of 8 and 16 seconds (Fig. 6B). In both cases, more surprising (e.g. less frequent) oddballs drove larger deviant responses (n=5). To investigate this effect further, we next designed sequences to compare responses between oddballs presented with equivalent frequencies but different element timing (Fig. 6C). We find that while deviant responses are generally larger for a 200 ms hold time than 400 ms, in both cases the OBS increases as oddball frequency decreases (Fig. 6D).

**Fig. 6.**
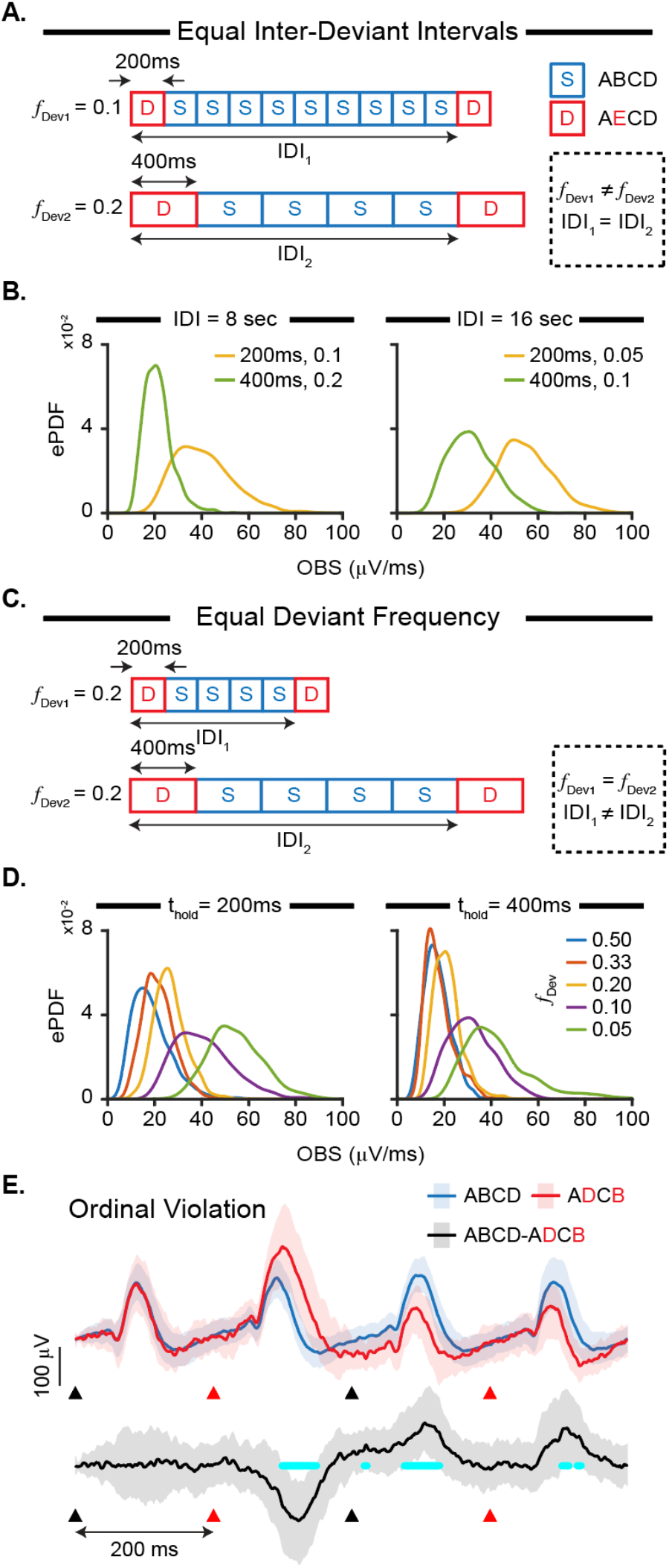
Simple neural adaptation cannot explain deviant responses. (A) Mice (n=5) were shown RGP sequences where the oddball presentation frequency, *f*Dev (0.1 when 1 out of 10 sequence presentations contains and oddball, 0.2 for 1 out of 5, etc.), and element hold times were varied to create a constant Inter-Deviant Intervals (IDI). (B) For both IDI = 8 and 16 sec, an Oddball Score (OBS) metric quantifying the deviant effect size as the time averaged difference between Std and Dev responses was larger for less common oddball events when thold = 200 ms than for more common events when thold = 400 ms sequences despite the fact that the IDI was the same. Plots show empirical probability distribution functions calculated from bootstrapped estimates of the OBS. (C) We next tested how the OBS would change as a function of IDI for constant *f*Dev values. (D) While the OBS was larger in sequences comprised of shorter elements, it increased as oddballs became less frequent (e.g. more surprising) for both hold times. Comparing (nested) linear models accounting for thold, *f*Dev, and IDI showed that oddball probability has a significant effect on the oddball score (*F*=320.66, *p* = 2.20e-62). E. We next used ordinal substitutions, in which the Dev sequence reorders elements in positions 2 and 4 rather than introducing a novel element, to control for both frequency and novelty effects. Reordered elements B and D result in significant deviant responses similar to those produced by novelty violations.

To confirm our impression that oddball stimulus probability has a significant impact on deviant responses, we used an F-test to compare fits produced by full and restricted nested linear models of our bootstrap OBS estimates. For the full model, we fit *OBS ∼ r + T + 1/P*, where *r* is presentation rate (e.g. 1/t_hold_), *T* is the time (in seconds) between deviant stimuli, and *P* is probability of seeing E given A. We then fit a restricted model that did not include a term for deviant stimulus probability. Both models were fit with 1000 data points (100 per condition). The data fit of the full model was significantly better than with the restricted model (*F* = 320.66, *p* = 2.22×10^-62^) indicating that oddball probability has a significant effect on the deviant response. We next repeated this analysis on directly computed oddball scores (i.e., non-bootstrapped, n=50) and the overall result was maintained (*F* = 4.59, *p* = 0.037). Since the probability of the oddball stimulus has a significant effect on the OBS, we conclude that deviant responses depend on both neuronal adaptation and stimulus likelihood.

Thus far we have only considered expectation violations created by introducing a novel element into a standard sequence. An alternative approach to drive prediction errors, that sidesteps novelty and more directly tests whether there is an explicit expectation of ordinal structure, is to substitute one familiar element with another. These ordinal violations result in sequence- and timing-specific alterations of neuronal activity over days (Gavornik and Bear, 2014; Price et al., 2023; Knudstrup et al., 2024a), and we sought to determine whether they also result in significant deviant responses in the single-session oddball paradigm.

To generate ordinal violations, mice (n=6) viewed ABCD 9 times followed by ADCB. The sequence ADCB contains an ordinal violation since image D is a member of the base sequence ABCD but is out of its usual order. As shown in Fig. 6E, replacing ABCD with the ordinal oddball ADCB created a large, significant deviant response to the 2^nd^-4^th^ elements of the sequence. In this sequence, the misplaced D occurs 200 ms after the D from the previous standard presentation (e.g. ABCDADCB) and is thus likely confounded by the short-term adaptation effect discussed previously. Indeed, the deviant response to the mis-ordered element D shows the same bias towards positivity seen in Fig. 5. This confound does not apply, however, to the 3^rd^ and 4^th^ elements since C is in its expected position within the sequence and B is outside the range where recency has a significant effect on VEP shape. The deviant responses in elements 3 and 4 are consistent with those seen in earlier experiments following an unexpected substitution oddball. From these findings, which conflict with findings in L2/3 (Homann et al., 2022), we conclude that ordinal expectation violations do generate significant deviant responses in V1.

### Deviant responses reflect conditional and higher-order sequential relationships

According to predictive processing and Bayesian theories of cortical function (Friston, 2005; Aitchison and Lengyel, 2017; Keller and Mrsic-Flogel, 2018), brain activity reflects a probabilistic model of the environment. In our previous experiments, the standard sequence has a fixed first-order probability, e.g. P(B|A) = 1. The overall probability across presentations, however, depends on the oddball frequency, e.g. P(B|A) = 0.9 when the oddball is present in 10% of presentations. As we have shown, deviant responses quantified with the OBS get larger as oddball frequency decreases which suggests an underlying representation accounting for relative event probabilities. Human experiments often use first-order conditional and second-order sequences in serial reaction time tasks, with performance depending on the ability of the test subject to recognize the relatively complex sequential structures. Though it seemed unlikely that V1 circuits would be capable of modulating oddball violations as a function of the conditional structure of these sequences, we decided to test this by presenting mice (n=12) with a 10-elelement first order conditional sequence borrowed from the human subject literature (DBDCABCADB, (Curran, 1997)) where element D was followed by both B and C, Fig. 7A. Though this sequence is ultimately deterministic, considering only the probability of direct transition from one element to another there is a 67% probability of transitioning to element B|D and 33% transition probability of C|D. To determine whether this makes a difference for deviant responses, we presented this sequence with an oddball element E following D at either a high or low probability location every 10^th^ presentation. Since there are three instances of element D, each mouse was run through three separate sessions, once with E replacing C, once with E replacing B in position 2 and once replacing B in position 10. In all cases, there was a large and significant deviant response to the oddball element. Despite the fact that our experimental design did not introduce any short-latency repeats of the oddball element (which in all cases was seen only once every 15 secs), the deviant response was biased in a positive-going direction. The magnitude of this response was noticeably larger and more persistent when the oddball occurred at one of the two high-probability locations (responses combined in Fig. 7A) than at the low probability location. This observation is confirmed by comparing the bootstrapped OBS distributions between these two cases which show that deviant responses are larger (Fig. 7B) when the oddball is presented to at one of the high-probability DB transition locations compared to the low-probability CB transition location.

**Fig. 7.**
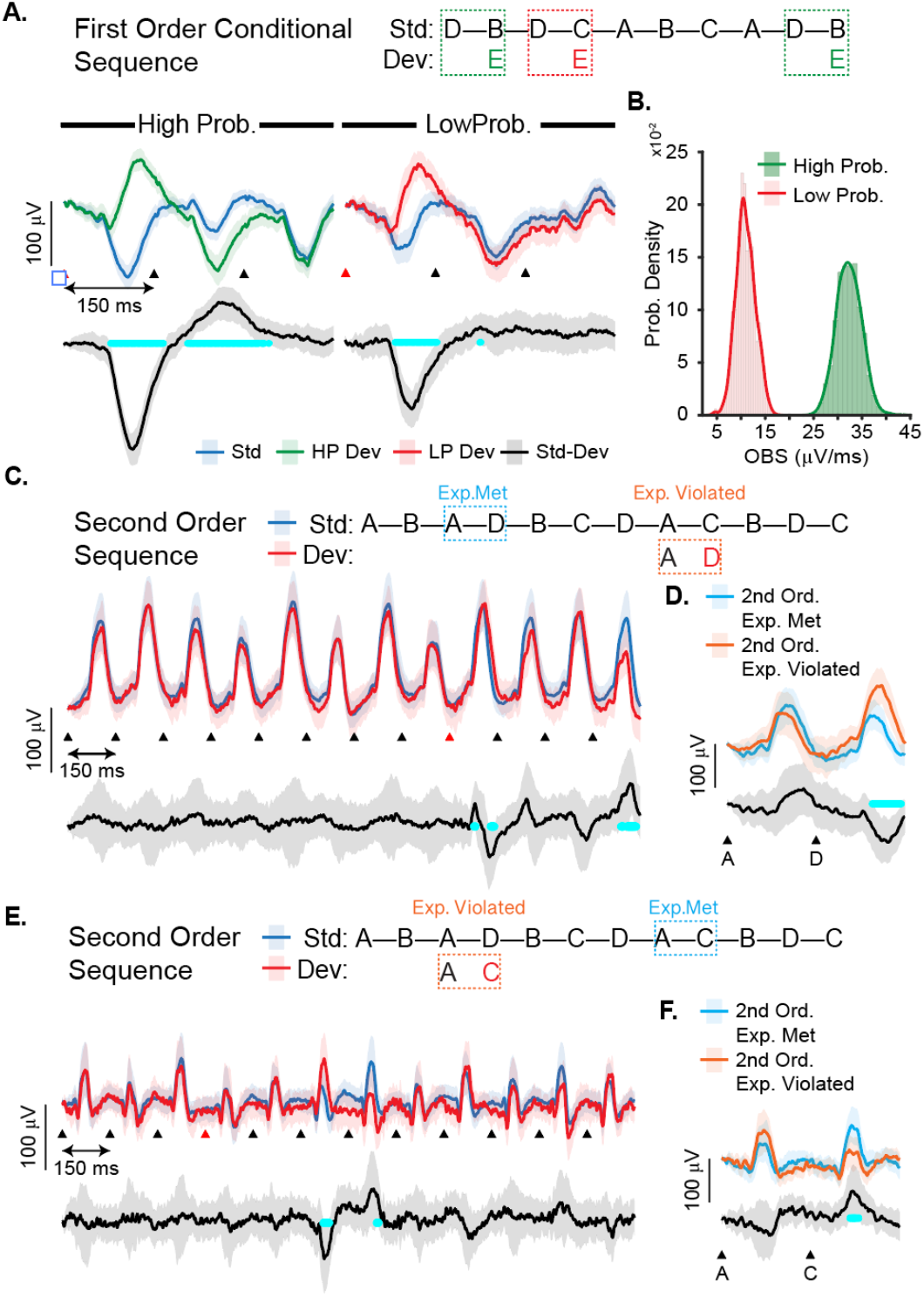
Deviant responses reflect conditional and second order relationships. (A) Mice (n=12) were shown first order conditional RPG sequences. In this sequence the first order probability of element B following D (High Prob) is ∼67% and C following D (Low Prob) is ∼33%. Significant deviant responses are seen when a novel element E follows D in any location (1 out of 10 sequences contained an oddball), though the magnitude and duration of the effect was larger when the substitution occurred in a High Prob transition location (left traces show the response averaged over both HP locations). (B) This observation is confirmed by plotting the distribution of bootstrap estimates of the oddball score (OBS) for both conditions. (C) To try and understand how the deviant response could scale based on position within the sequence, we showed mice (n=13) a 12-element second order sequence containing each possible first-order transition from A (AB, AC, and AD). Dev sequences containing the valid first-order transition AD at the location of the AC transition drove a significant difference relative to Std sequences immediately after the second order violation. (D) The response to element pairing AD was significantly different when the second order expectation was violated compared to the response produced by the same element pairing when the expectation was met (locations within the sequence marked in panel D, 2^nd^-order statistics are calculated at the 97.5% confidence intervals to control for multiple comparisons). (E) Element D presentations were separated by a single element in the deviant sequence, raising the possibility the violation signal was confounded by the recency effect (see Fig. 5). To control for this, the same experiment was repeated on a second cohort of mice (n=8) using the same sequence but with C replacing the D in position 4. This substitution, representing a valid first order transition, did not drive an oddball response at the time of the substitution but did result in a significant difference following the C-B-C sequence that we attribute to the recency effect. (F). The response to A-C is significantly different when it violates second order expectations (B-A-C) compared to the expected (D-A-C).

These results comport broadly with the previous demonstration that deviant responses scale with surprise, loosely defined as the inverse of the oddball occurrence frequency. In this interpretation, it is less surprising when an oddball is more predictable because it occurs more frequently (e.g. *f*_Dev_ = 0.2, or 1 oddball for every 5 sequence presentations) than less (e.g. *f*_Dev_ = 0.1, or 1 oddball for every 10 sequences). Deviant responses here were larger when they occurred in high-probability event locations, again suggesting that more surprising events generate larger neural responses. While the first order sequence is, as mentioned above, fully deterministic its structure does create second order ambiguity since BD can lead to either a B or a C. To look more carefully at the extent to which second order structure modulates visual responses, we next ran a group of mice (n=13) using a 12-element second order sequence previously used with human subjects (ABADBCDACBDC, (Curran, 1997)). This sequence contains every possible first order transition from element A (e.g. P(B|A) = P(C|A) = P(D|A) = 1/3) but these can be uniquely distinguished from each other based on second order structure (BAD, DAC, and CAB). We tested whether substitution of a valid first order transition that violated this second order structure would produce a deviant response by replacing the AC transition with AD in every 10^th^ sequence presentation. As shown in Fig. 7C, this substitution did result in a large and significant deviance response at the time of the substitution. There was also a significant difference in the response to AD when it occurred at the expected location compared to the unexpected location (Fig. 7D). While this finding was intriguing, we realized that our oddball structure introduced a DAD transition sequence potentially confounding the deviant signal based on the recency effect. To correct this, we ran a new cohort of mice (n=8) with a reordered the sequence (ABADBCDACBDC, Fig. 7E) and switched the location of the oddball to compare AC transition responses when they met and violated second order structure. In this new sequence, previous presentations of C before AC in both the expected and unexpected locations occur several elements in the past to minimize the recency effect. In this new experiment, the oddball no longer generates a significant deviant response at the time of the substitution (though it does create a significant deviance for the following C presentation which occurs within the adaptation window) but there is a significant difference between AC when it is seen in its expected and unexpected locations (Fig. 7F). Taken together, these results are consistent with the hypothesis that visual experience is sufficient to entrain weak representations of second order expectations that can modulate visual responses when valid first-order transitions violate second order structure.

## Discussion

The oddball paradigm is a well-established tool used to study cortical responses to sensory signals in different contexts. An ongoing question is the extent to which MMN, which has been reported in both humans and various animal models, reflects a neural signature of deviance detection or is a consequence of relatively simple stimulus-specific adaptation (Ross and Hamm, 2020). Whether or not MMN represents evidence of predictive coding has been widely considered (Wacongne et al., 2012; Stefanics et al., 2015, 2016; Wacongne, 2016; Gallimore et al., 2023), but the answer is still unclear. If oddball-evoked deviants do represent evidence of predictive coding, it seems reasonable that deviance signals would appear when sensory inputs violate expectations that are more complex than novelty based on event frequency. Our findings suggest that this does occur. Specifically, we show that VEPs calculated from the LFP clearly generate differential responses to unexpected visual stimuli across a wide range of times, that deviance signals and multi-day plasticity is influenced by the temporal structure and regularity of visual sequences, that the size of the deviant responses increases as the oddball becomes less predictable, and even that higher order expectation violations drive differential responses.

Most experiments studying MMN, or employing it for clinical purposes, use a simple design where the oddball stimulus occasionally interrupts a repetitive sequence of a single stimulus (e.g. A – A – A – A – A – B – A – A …) (Pazo-Alvarez et al., 2003; Kujala et al., 2007; Kremláček et al., 2016) with responses recorded during a randomized “many standards” protocol used as a control (Ross and Hamm, 2020). More directly relevant to this work are two recent studies that used more complex sequential structures and stimulus elements study deviance signals in mouse V1 by measuring activity dependent Ca signals. The first focused on characterizing excess activity in layer 2/3 neurons evoked by a novel oddball inserted into regularly repeating sequences (Homann et al., 2022). This study was notable for introducing the RPG stimulus and concluded that excess activity generated by novel elements can be explained by an adaptive accommodation model based on “generic, widely available mechanisms”. Key to this conclusion was their finding that re-ordered sequence elements do not create significant excess activity. In our experiments, by contrast, re-ordering drove a large highly significant deviance signal (Fig. 6E).

One possible explanation for this discrepancy is that LFP-based VEP reports average summated synaptic currents across cortical layers and dendritic segments that may not result in obvious changes in population-level Ca^2+^ traces in L2/3 excitatory cells (Halnes et al., 2024). It is also important to note that their conclusions and model were based on a determination that reordered sequences drove “very little excess activity”, defined specifically as increased fluorescence, compared to novel elements. However, a careful examination of their results (Homann et al. 2022, Fig. 4B) shows that reordered elements do drive a small, and apparently robust, differential response characterized by an initial decrease in DF/F signal followed by a period of increase DF/F lasting approximately 2 seconds. This signal, which is not analyzed as it does not meet their definition of excess activity, could be the somatic Ca reflection of the LFP-based signal recorded in our experiments. In any event, it is hard to reconcile their synaptic adaptation model, which is based on exponential decay relative to stimulus times, with our findings that deviant response magnitude scale as a function of oddball frequency for fixed inter-deviant-intervals (Fig. 6B), sequence order (Fig. 6D), or more complex statistics within a sequence structure (Fig. 7).

The second study examined how Ca^2+^signals in different layers of V1 respond to pattern-violating images over multiple days of visual exposure (Gillon et al., 2024). This work from the Allen Institute imaged from dendrites and soma in different cortical layers across days using RPG-style sequence elements but with a more abstract definition of novelty. Familiar sequences consisted of four unique frames of randomly placed Gabors, each rotated to the same orientation. A pattern violating image would differ from the preceding frames in both the position and rotation of its constituent Gabor patches and all sequences were separated by a gray frame (which we found is important for driving robust plasticity, Fig. 2). While they found large, significant differences in Ca^2+^ responses driven by pattern violating images these did not emerge as statistically significant until the 3^rd^ day of imaging in layers 2/3 and 5. There are enough methodological differences in experiment design that it is hard to compare our results directly with theirs, but we are in general agreement with their conclusion that V1 deviant responses are consistent with some aspects of predictive coding models. It would be very interesting to examine whether our stimulus paradigms drive differential deviance signals across layers similar to those reported in this study.

Unlike in previous multi-day sequence plasticity experiments (Gavornik and Bear, 2014; Sidorov et al., 2020; Finnie et al., 2021; Price et al., 2023; Knudstrup et al., 2024a), we find no evidence that V1 actively predicts the specific timing of elements in the standard sequence. In these studies, multi-day exposure to OSG sequences drove VEP potentiation that is selective for stimulus timing (Gavornik and Bear, 2014) and causes L4 spiking and L2/3 Ca^2+^ signals to increase when an expected element is omitted (Price et al., 2023; Knudstrup et al., 2024a). Comparing our work here with previous findings suggests that it takes more time for the brain to generate a model of when an expected event will occur, and it may be that the mechanisms responsible for generating in-session deviant responses in V1 are not as complex as those engaged during a multi-day sequence learning task. We were surprised that the inclusion of a gray screen between sequence presentations has such a large effect on multi-day plasticity since activity evoked during active stimulation is presumably responsible for driving plasticity and the same in Figure 2 panels. We have previously shown that multi-day sequence plasticity requires muscarinic acetylcholine receptors (Sarkar et al., 2024), and also that ACh release is retinotopically mapped by visual stimulation (Knudstrup et al., 2024b), which might suggest a large synchronizing ACh release event occasioned by transitions from gray to patterned stimuli is required to activate underlying plasticity mechanisms. Relatedly, while a functional hippocampus is required for V1 sequence plasticity (Finnie et al., 2021), we are unaware of any similar hippocampal requirement for MMN style deviant responses. Finally, while multiple studies have found that NMDA receptor are required for MMN signals, antagonizing these receptors does not prevent sequence plasticity (Gavornik and Bear, 2014). Overall, these observations demonstrate that cortical circuits recognize spatiotemporal and ordinal structure using different mechanisms operating at different timescales. Future work clarifying how these relate to each other will be important to inform predictive coding models, and specifically the plasticity rules required to create predictive models of sensory experience.

In summary, our findings suggest that visual circuits develop relatively complex expectations of sequential structure at a rapid timescale and generate responses that differentiate between expected and unexpected visual transitions. This is generally consistent with a predictive coding model of cortical function (Keller and Mrsic-Flogel, 2018), though many details remain to be worked out as they relate to the underlying mechanisms. While previous work dissecting visual MMN circuits in simpler oddball paradigms (Hamm and Yuste, 2016; Hamm et al., 2021), in more abstract sequences (Gillon et al., 2024), or in behavioral tasks including visual flow and motor afferents during locomotion (Keller et al., 2012; Leinweber et al., 2017), provide relevant insight more work is required to explain our findings and weave the various experimental threads into a single unified theory of cortical responsiveness and predictive coding.

## Acknowledgements

We thank Byron Price for his advice and consultation on experiment design and data analysis. This work was supported by the NEI (R001EY030200) and NIMH (R00MH099654).

## Author Contributions

Conception and experimental design: J.P.G., S.K, R.S. Surgeries, experiments, and data collection: S.K., C.R., C.M.J., R.S., and M.F. Data analysis: S.K., C.R., J.P.G. Writing and figures: S.K. and J.P.G.

